# Neocortical microdissection at columnar and laminar resolution for molecular interrogation

**DOI:** 10.1101/200931

**Authors:** Koen Kole, Tansu Celikel

## Abstract

The heterogeneous organization of the mammalian neocortex poses a challenge to elucidate the molecular mechanisms underlying its physiological processes. Although high-throughput molecular methods are increasingly deployed in neuroscience, their anatomical specificity is often lacking. Here we introduce a targeted microdissection technique that enables extraction of high-quality RNA and proteins at high anatomical resolution from acutely prepared brain slices. We exemplify its utility by isolating single cortical columns and laminae from the mouse primary somatosensory (barrel) cortex. Tissues can be isolated from living slices in minutes, and the extracted RNA and protein are of sufficient quantity and quality to be used for RNA-sequencing and mass spectrometry. This technique will help to increase the anatomical specificity of molecular studies of the neocortex, and the brain in general as it is applicable to any brain structure that can be identified using optical landmarks in living slices.

## Introduction

The mammalian brain displays a high degree of cellular and anatomical heterogeneity, with specific distributions of numerous cell classes within individual brain regions. The neocortex, for example, is organized in a laminar manner, each layer with distinct functional roles, abundances of cell-types (neuronal or non-neuronal) and gene expression profiles ^1–4^. Such heterogeneity might allow functional diversity required for complex computations leading to perception, learning and memory. However, it also introduces a challenge for researchers aiming to disentangle the molecular mechanisms of network organization and function.

High-throughput methods (e.g. (epi)genomics, transcriptomics, proteomics, metabolomics) are becoming increasingly well-established and accessible, allowing for large-scale interrogation of molecular changes in the brain. Ideal application of -omics screening requires identification of molecular profiles of cells and populations in their network environment. However, current methods allow either isolation of individual cells/cell types ^5–7^ or extraction of large tissue sections aggregating cells across anatomical (sub-)regions (e.g. columns and layers) across several cubic millimeters of brain tissue ^8–10^.

In the neocortex, feasibility of laminar tissue isolation has been previously shown ^11^, however due to the columnar nature of cortical processing, molecular mapping of network organization and function requires sample isolation at both columnar and laminar resolution. Here we introduce such a technique that allows isolation of very small tissue sections (~350 μm) from acutely prepared brain slices. The method is based on visually guided microdissection in brain slices, and is an expansion of the acute electrophysiological slice preparation, commonly used in the central nervous system, e.g.^12–16^.

### Comparison with other methods

While single-cell RNA sequencing can be used to study transcriptomic changes of individual cells in isolation ^5,6^, analysis of whole-tissue samples remain the most commonly used approach in -omics research. Obtaining these samples at high anatomical resolution is, however, challenging. Both RNA and proteins are prone to degradation, an issue commonly combated by keeping samples frozen (**Table 1**). However, tissues tend to crack when they are too cold which interferes with the subsequent identification of region of interest (ROI) and its isolation. Although neuroanatomical landmarks can be used to identify the ROI, some structures, such as supragranular, infragranular, and granular cortical laminae, or the septa that separates cortical columns in most granular cortices, cannot be reliably distinguished in frozen tissue. Specific imaging conditions (e.g. infrared DIC microscopy ^17^, Dodt gradient contrast ^18^, spectroscopic imaging of intrinsic markers ^19^) might be required for ROI visualization before cryopreservation. A method that allows sample isolation in (near) physiological states of the tissue with high anatomical accuracy, while preserving sample quality, is thus required for molecular interrogation of anatomically and functionally diverse regions of the brain.

**Table 1.** A comparison of tissue isolation methods for molecular biological analysis. Fields marked in bold represent the most favorable experimental conditions/outcomes.

| *Dissection method* | *Specialized infrastructure requirement* | *Anatomical resolution* | *RNA and protein quality* | *Labor intensity* | *Intermediate processing required after slice experiments* | *Citations* |
| --- | --- | --- | --- | --- | --- | --- |
| Whole tissue punch from frozen tissue | **No** | + | +++ | ++ | Yes | 8–10,21,22 |
| Micro tissue punch from frozen tissue | Yes | ++ | +++ | + | Yes | 23 |
| Laser capture microdissection | Yes | +++ | + | +++ | Yes | 20,24–27 |
| Whole tissue punch or macrodissection from acute slices | **No** | + | +++ | ++ | **No** | 15,28,29 |
| Acute brain slice microdissection | **No** | ++ | +++ | ++ | **No** | Kole et al, 2017a<br>Kole et al, 2017c |

Acutely prepared brain slices are indispensable for *ex vivo* electrophysiological experiments in neuroscience. Due to the widespread use of acute slices, the method described herein can be established in many (if not most) cellular/molecular neurobiology and synaptic/network physiology labs without additional infrastructural investment and with simple modifications in the pipeline (**Table 1**). Obtaining microsections from acute brain slices, as opposed to frozen or fixed tissues, as required by laser-guided microdissection ^20^, has the advantage that samples can be collected directly after acute manipulations, e.g. induction of synaptic plasticity, incubation with different pharmacological substances, after cellular stress, among others, without the need for intermediate freezing, fixation or cryosectioning (**Table 1**). An important consideration however is the timespan in which tissues are collected. Cells in acutely prepared brain slices undergo transcriptional and translational processes that might affect molecular profiles. Thus, processing times should be minimized, and kept as similar as possible across experimental groups.

During *Acute Brain Slice Microdissection*, the method described here, ROIs are extracted sequentially. This step-wise tissue excision (as opposed to the one-shot approach with tissue punches) has the advantages of (1) allowing potential rectification of mistakes during excision, (2) excision of brain tissues that are situated (very) closely, even next to one another, (3) excision of samples in a non-circular manner (tissue punchers are typically round), conforming to the anatomical boundaries of the ROI.

### Application of the method

We have used the protocol described herein to identify the transcriptomic and proteomic changes associated with experience dependent plasticity in single column and laminar resolution in the mouse somatosensory cortex ^30–33^. Samples collected using this method could however potentially be used for any -omics screening, in any brain region that can be identified in acute brain slices. Using region-specific promoters to drive fluorophore expression (e.g. ^34–37^), viral expression of fluorophores in anatomically targeted locations ^12,38,39^ or retrograde vectors to identify projection neurons ^40^ ROI selections could be extended to arbitrary regions defined anatomically, functionally, cellularly or based on their connectivity profiles. Although we thus far processed microdissected tissues in their entirety, it might be possible to introduce a cell sorting step between tissue isolation and downstream processing as the approach described herein does not require snap freezing of the sample prior to isolation. Cell sorting would increase the anatomical resolution further and would provide insight into laminar specificity of gene expression within and between cell types. However, it would require pooling samples across different slices and animals. Because the acute slicing procedure we have adapted is commonly used for *in vitro* electrophysiological and pharmacological experiments, various physiological, biochemical and pharmacological manipulations could potentially be employed to further expand the utility of the -omics screening in systems neurobiology.

### Limitations of the current technique

Due to ongoing cellular processes, the tissue collection requires rapid handling times. In comparison to tissue extraction after cryosectioning, which allows slices to be stored for extended periods of time before tissue collection, our approach will require pilot experimentation by the novice experimenter to minimize handling times. The microdissection, just like manual tissue punching, requires a steady hand from the experimenter; it is advisable to initially practice the technique with dummy samples. We have experimented with micromanipulators to manoeuvre the microdissection needle, however in our hands this greatly increases handling time and does not allow utilization of the tactile feedback present during manual dissection. In our opinion, manual manoeuvring of the needle is thus more favorable.

### Experimental design and considerations

Depending on the experimental goals and the ROI, varying amounts of material can be collected on a single day (**Figure 1**). In our experience, if only tissue excision is required without any further manipulations (e.g. electrical stimulation, plasticity induction, pharmacological incubation), tissues from ~5 animals can be obtained on one day. This does however require careful planning, as tissues should be collected as soon as possible following slice preparation. The quality and content of extracted materials (RNA, protein, metabolites) depend heavily on the viability of the slices. Fresh slices typically contain the highest number of healthy cells. With cell degradation over time *ex vivo*, changes in transcription and translation will be reflected in the tissue extract at the molecular level.

**Figure 1.**
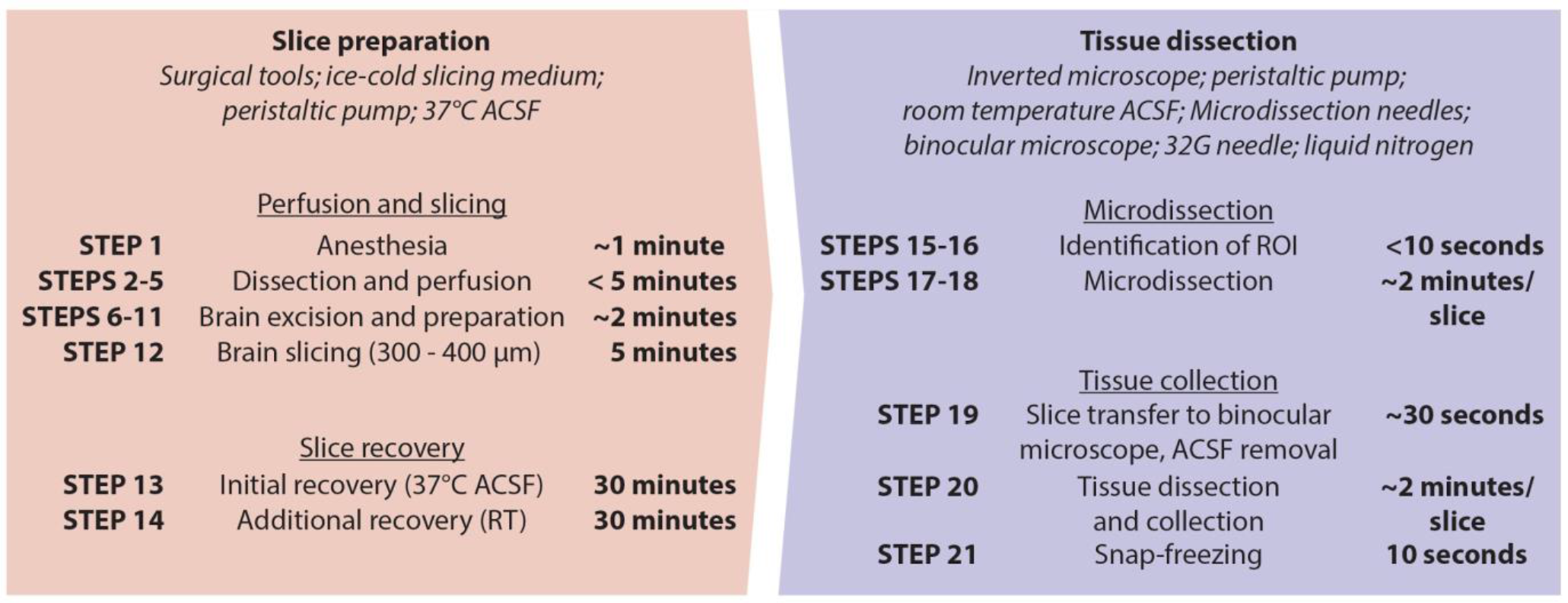
Flowchart of slice preparation, microdissection and tissue collection. Essential equipment and media are listed *in italics*. From anesthesia to end of slice recovery, slice preparation takes ~1 hour to complete, while tissue collection can be completed within 5 minutes per slice depending on complexity of the required excision. Care should be taken to reduce the time until tissue collection.

We typically prepare slices from two animals consecutively, with the tissue extraction of the first animal overlapping with recovery of the slices from the second (see **Figure 1** for the experimental stages). After tissues have been collected from the first two animals, additional slices are prepared. Hence, changes in transcription and translation over time, which are typically not of interest and could introduce variance between samples, are kept to a minimum. If other experimental manipulations are a part of the experimental design, such as electrophysiological, optogenetic, pharmacological interventions, the timing of slice preparation should be adjusted (**Figure 1**). Since microsections yield low amounts of material, it may be necessary to pool tissues. Because typically more than one slice and hemisphere can be used, depending on the experimental design, it is possible to pool tissues from the same animal. Before using samples for any downstream application, it is advisable to first use a small set of pilot samples to ensure sufficiency of quality, quantity and purity of the obtained samples (see *Anticipated results* below).

For our experiments, we required an upright microscope with Dodt gradient illumination in order to visualize barrels in L4 of the primary somatosensory cortex, barrel cortex subfield (**Figure 2B**). Although such a setup might not be required to visualize other brain regions (e.g hippocampus, olfactory bulb, striatum), visualization of the microdissection needle is enhanced in an upright microscope (as opposed to an inverted microscope), increasing the precision and hence reproducibility of tissue microdissection. Further enhancement can be achieved with the use of the appropriate microdissection needle diameter and holding angle (**Figure 2D** and **E**).

**Figure 2.**
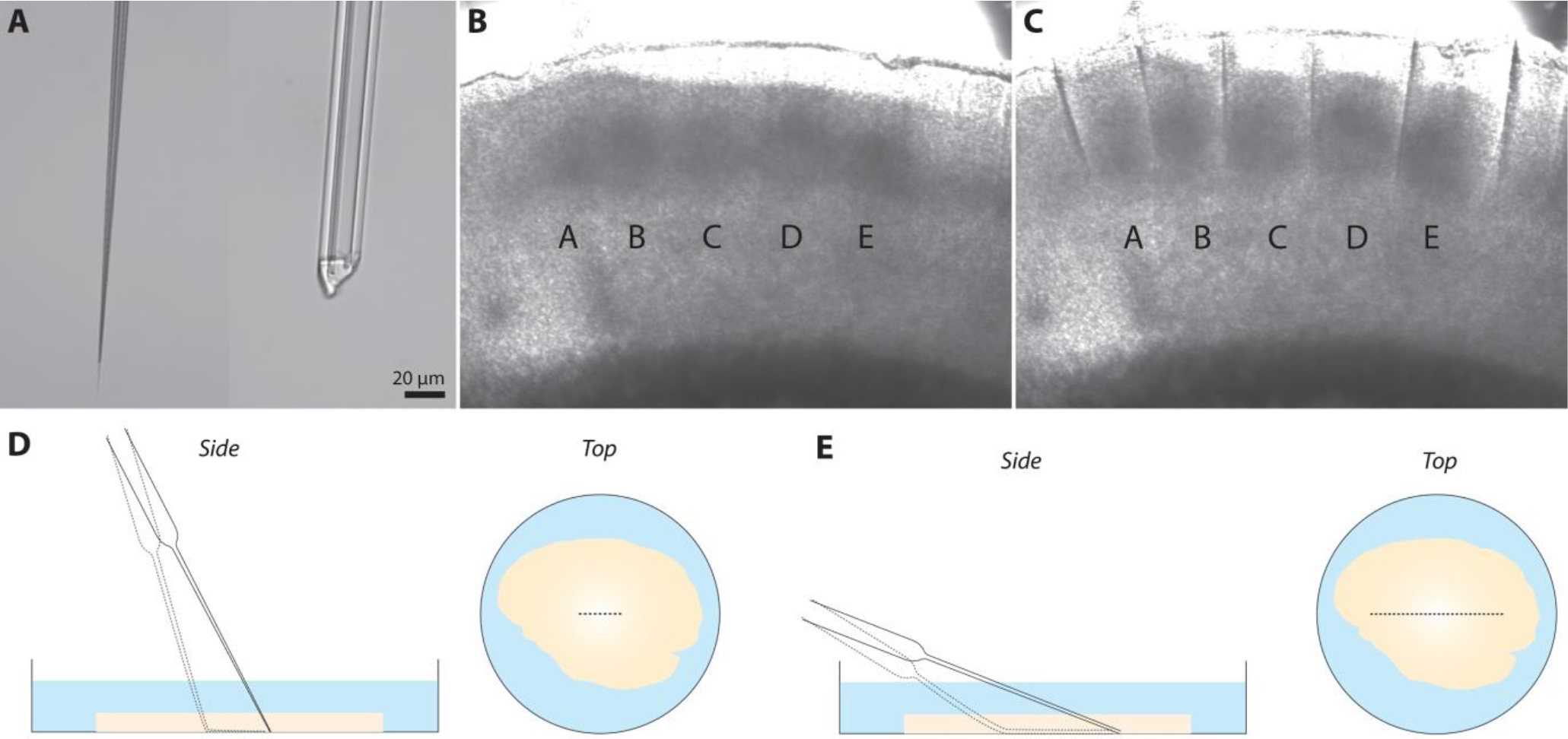
Microdissection of single cortical columns in the mouse somatosensory cortex. **(A)** Glass microdissection needles at high magnification. Left, glass pipette immediately after pulling; Right, the same pipette after controlled tip trimming. A shorter tip with a sharp edge results in reduced bending moment, increased stability and ease of incision. **(B)** Bright-field image of acute 300 μm mouse thalamocortical slice. Cortical columns A-E can be identified by barrel structures in L4. **(C)** A dissection needle as shown in **A** (right) was used to create intercolumnar cuts. These can subsequently used to identify individual columns by eye under a dissection microscope. **(D)** An uncut needle is more flexible at the tip. Hence it can be bent at a steeper angle, making it suitable to make shorter incisions. Note that at lower angles, the pressing force at the tip may not be sufficient to make visible incisions. **(E)** A microdissection needle cut to a ~20 μm diameter has reduced flexibility at the tip. Held at a low angle, needles such as these can be used to make long incisions. The reduced tip flexibility ensures sufficient pressure at the tip, but simultaneously prohibits strong bending.

Finally, if the experimental design requires targeted alteration of molecular/cellular function, upon e.g. sensory deprivation ^14,41^, induction of synaptic plasticity in identified pathways ^13,42^, systematic modulation of cellular excitability ^15,16,43^, circuit alterations ^38,39,44–46^ or behavioral training ^44,47,48^, it would be advisable to design the experiments to ensure availability of within animal, and within slice (wherever applicable), controls to minimize the genomic variability across independent samples and the experimental groups.

## Materials

### Reagents

#### Media preparation

- Sterile Milli-Q water (18 MOhm), e.g. autoclaved
- Choline chloride (ChCl; Sigma #7527-500G) ***CRITICAL*** ChCl should be stored under inert gas (e.g. nitrogen)
- Sodium chloride (NaCl; Merck #1064041000)
- Sodium bicarbonate (NaHCO_3_; Merck #1063291000)
- Sodium phosphate monohydrate (NaH_2_PO_4_.H_2_O; Merck #106346500)
- Potassium chloride (KCl; Merck #1049360500)
- Glucose monohydrate (C_6_H_12_O_6_.H_2_O; Merck #1083422500)
- Calcium chloride dihydrate (CaCl_2_.H_2_O; Merck #1023820500)
- Magnesium sulphate heptahydrate (MgSO_4_.7H_2_O; Merck #1058860500)
- Sodium pyruvate (Na-Pyruvate; Sigma #P2256-25G) ***CRITICAL*** Na-Pyruvate should be stored at 4 degrees Celsius

#### Slice preparation and recovery

- Experimental animals ***CAUTION*** Use of experimental animals must be in accordance to national regulations
- Carbogen (95% O_2_, 5% CO_2_)
- Isoflurane
- Cyanoacrylate Glue
- Ice
- Agarose (2%) (optional)

#### Tissue dissection

- RNAseZap (Sigma #R2020)
- Liquid nitrogen (N_2_)
- Dry ice

#### RNA isolation

- miRNeasy Mini Kit (Qiagen #217004), or other RNA isolation method of choice

### Equipment

#### Perfusion and slice preparation

- Sterilized surgical tools

- Large scissors
- Large tweezers
- Small scissors
- Fine tweezers
- Clamp
- Small scoop
- Large scoop
- Microscope slide
- Razor blade
- Peristaltic pump (for mice: 6 ml/minute flow rate for perfusion), tube exit fitted with a needle (23 Gauge) A 20 ml syringe filled with SM and fitted with a needle can also be used
- If agarose embedding is used: Water bath, temperature set to keep agarose in a liquid state
- Water bath, 37 degrees Celsius
- Water bath, room temperature
- Anesthesia box: add ~0.5 ml of isoflurane to a small box or bell jar for initial anesthesia
- Anesthesia ‘face mask’: add ~0.5 ml of isoflurane to a 50 ml tube with a ball of paper tissue
- Vibratome, e.g. Precisionary Instruments VF-300
- Air stones or other means to carbogenate media
- Beaker or other container to hold the brain after removal from the skull
- Slice storage chambers (e.g. Automate Scientific #S-BSK4 or can be custom made out of rings cut from a 10 ml syringe, and three strings from a nylon hose glued to its rim in non-orthogonal angles using cyanoacyrlate)

#### Tissue (micro)dissection

- Pipet puller, e.g. Sutter Instruments P2000
- Borosilicate glass capillaries, e.g. Harvard Apparatus 30-0034, OD 1.0 mm, ID 0.50 mm
- Upright microscope with camera and appropriate lighting source to discern ROI
- Holding chamber with sufficient space to manoeuvre the dissection needle; In our experience the lid of a 35 mm petri dish is well suited (e.g. ThermoFisher #153066) ***CRITICAL*** A platform should be present next to the holding chamber on which the experimenter can rest their hand during microdissection. If absent, the incision precision might suffer due to lack of mobile stability.
- Peristaltic pump (1 ml/minute flow rate)
- Binocular microscope
- Flexible dissection mat
- Insulin needle, 32G or smaller
- Sterile Eppendorf tubes
- A slice anchor that keeps the tissue in place, (e.g. Warner Instruments #64-0254, or can be custom made from a polymer ring with two strings of fine dental floss attached to its rim with a non-orthogonal angle).

#### RNA isolation

- Table-top centrifuge
- Sterile, RNAse-free Eppendorf tubes
- Sterile, RNAse-free pipette tips
- Depending on the expected RNA quantity: Qubit and necessary reagents (<40 ng/μl; e.g. Thermo Fisher #Q33217) or Nanodrop Spectrophotometer (>40 ng/μl; e.g. Thermo Fisher #ND-2000)

#### RNA integrity control

- Agilent BioAnalyzer or TapeStation
- In case of low quantities: BioAnalzyer Pico Kit (<5 ng/μl) (Agilent #5067-1513) or TapeStation High Sensitivity RNA Screentape and sample buffer (<10 ng/μl) (Agilent #5067-5581 and #5067-5580)
- In case of medium to high quantities (>25 ng/μl): BioAnalyzer Nano Kit (Agilent #5067-1511) or TapeStation RNA Screentape and sample buffer (Agilent #5067-5576 and #5067-5577)

### Reagent setup

#### Slice preparation and recovery

- Slicing medium (SM; 108 mM ChCl, 3 mM KCl, 26 mM NaHCO_3_, 1.25 mM NaH_2_PO_4_, 25 mM glucose, 1 mM CalCl_2_, 6 mM MgSO_4_ and 3 mM Na-pyruvate, or other recipe of choice that yields physiologically viable slices) ***CRITICAL*** SM must be prepared fresh every day ***CRITICAL*** SM must be carbogenated for 30+ minutes before use ***CRITICAL*** SM must continuously be carbogenated throughout use ***CRITICAL*** SM must be ice-cold (2-4 degrees Celsius) throughout use
- Artificial cerebrospinal fluid (ACSF; 120 mM NaCl, 3.5 mM KCl, 10 mM glucose, 2.5 mM CaCl_2_, 1.3 mM MgSO_4_, 25 mM NaHCO_3_ and 1.25 mM NaH_2_PO_4_) ***CRITICAL*** ACSF must be prepared fresh every day ***CRITICAL*** ACSF must be carbogenated for at least 30 minutes before use ***CRITICAL*** ACSF must be continuously carbogenated throughout use ***CRITICAL*** ACSF used for slice recovery should be at 37 degrees Celsius before use (tissue dissection can be done at room temperature)

### Equipment setup

#### Perfusion and slice preparation

- All surgical tools, the microscope slide and razor blade should be clean and sterile. The tools and working area can be disinfected using 70% ethanol
- Place a container on ice and fill it with ~250 ml SM (ensure carbogenation). This will allow gentle removal of the brain from the skull and cools down the brain.
- Optional (when using a Precisionary VF-300 Microtome or similar vibratome): Prepare 2% agarose and keep it in a liquid state using a water bath.

#### Tissue dissection

- Pipet puller settings: to obtain pipettes such as in **Figure 2A**, we used the following settings with our P2000 pipette puller:
  - Heat: 400
  - Filament: 3
  - Velocity: 35
  - Delay: 180
  - Pull: 1000
  The aim is to obtain long (~1 cm), thin, flexible pipette tips, to be used as microdissection needles. Sterile scissors or a razor blade can be used to cut the distal tip of the pipette to obtain a tip such as in **Figure 2B**. These stocky tips are suitable for tissue excision from cortex (**Figure 2C** and **D**); When other brain regions are of interest, pipettes of different diameters and lengths might be required
- The upright and binocular microscopes should be as close to one another as possible to ensure a smooth workflow.
- The screen displaying the camera image should be placed so that the experimenter can easily manoeuvre their microdissection needle while simultaneously viewing their actions.
- If RNA is to be extracted, clean microdissection needles, insulin needle and dissection mat with RNAseZap to reduce introduction of RNAses into the sample.
- Perfuse the holding chamber under the microscope with room temperature carbogenated ACSF at 1 ml/min using a peristaltic pump.

#### RNA isolation

- Prepare the solutions of the RNA isolation kit, e.g. miRNeasy Mini Kit
- Ensure the working area, pipettes, gloves, pipette tips are RNAse-free. Use RNAseZap to remove RNAses from the working area and equipment.
- Keep all samples on dry ice until lysing tissue in lysis reagent (such as Trizol or QIAzol)

## Procedure

### Perfusion and preparation of acute brain slices

***TIMING*** 10 - 15 minutes

***CAUTION*** Ensure proper ventilation to avoid inhalation of isoflurane

***CAUTION*** Biological waste (cadavers) should be disposed of according to institutional guidelines

***CRITICAL*** Work quickly but gently to avoid damaging the brain. Avoid the brain being exposed to air by keeping it submerged in, otherwise keep it moist using, SM. Avoid contact with the brain as much as possible

***CRITICAL*** Perfusion of the brain is required to remove blood from the tissue, which could affect downstream -omics profiles

#### TROUBLESHOOTING

1. Anesthetize the animal in the anesthesia box using isoflurane. After the animal reaches stage III-4 of the anesthesia ^49^ as determined by the lack of vibrissae movement, pinch withdrawal, corneal reflex and eyelid reflex, move the animal to the dissection area and place the head in the anesthesia mask. ***CAUTION*** Ensure full anesthesia before proceeding dissection. Extend anesthesia if necessary.
2. Using large scissors and large tweezers, open the abdominal cavity and remove the skin covering the chest. Cut away the diaphragm, ribs and sternum to open up the chest cavity and allow free access to the heart.
3. Clamp the aorta caudally to the diaphragm. This ensures swift replacement of blood in the brain by slicing medium during the perfusion.
4. Place the needle of the peristaltic pump in the left atrium, being careful not to damage the heart. Cut the right atrium using fine scissors and immediately increase pump speed to 6 ml/minute. Perfuse the animal for 1 to 2 minutes.
5. Remove the needle and turn off the pump. Use large scissors to decapitate the animal.
6. Use fine scissors to cut open the skin over the skull.
7. Cut open the skull with a second set of clean fine scissors to avoid infection with bacteria from the animal's skin and fur. Make two lateral incisions (left and right) from the caudal part of the skull. Make one incision along the entire midline.
8. Using tweezers, fold open the skull on the left and right side, revealing the brain. ***CRITICAL*** Be careful not to stretch the dura, which could cut into the brain, damaging it. This is critical for cortical experiments.
9. Using a small scoop, remove the brain from the skull. Gently transfer the brain from the skull directly into the container with SM.
10. Scoop up the brain with a large scoop and place it on the microscope slide or another clean, flat surface. Use the razor blade to quickly remove the cerebellum (in the case of coronal slices of cerebrum).
11. Glue the brain onto the appropriate vibratome cutting stage. Optional (when using a Precisionary VF-300 Microtome or similar vibratome) embed brain in 2% agarose. Quickly cool down the agarose to solidify it.
12. Transfer the stage and the brain to the vibratome, submerge it in SM and cut 300-400 μm slices. ***CRITICAL*** Depending on the age of the animal, different thicknesses are more suitable. Brains of juvenile and young adult animals tend to be softer and less rigid, and hence benefit from thicker (350-400 μm) slices; conversely, adult animals may require thinner (300-350μm) slices to allow nutrients to diffuse through the denser tissue.

#### TROUBLESHOOTING

13 Gently transfer each slice to brain slice storage chambers in the 37 degrees Celsius ACSF.Leave slices to recover for 30 minutes.

14 After initial recovery, allow ACSF to return to room temperature. This may be sped up by transferring the ACSF container to a water bath at room temperature water.

If samples need to be extracted from multiple animals on the same day, the recovery periods might be used to prepare slices of the next animal.

### Tissue dissection

***TIMING*** Approximately 5 minutes per slice depending on the complexity of ROI to be excised

15 Select a brain slice containing the ROI and transfer it to the holding chamber. Place the slice anchor, ensuring that the ROI is flattened and the tissue cannot move during incisions.

***CRITICAL*** Proper placement of the slice in the holding chamber is important to ensure ease of access during microdissection. For instance, if the microdissection needle approaches the slice from the right side and cortical tissues are to be dissected, face the cortex towards the right.

16 Using an objective lens that allows a large field of view (we typically use 4x), locate the ROI

17 While using the magnified view, gently approach the brain slice with a microdissection needle, holding it between thumb and index finger while resting the hand next to the holding chamber. Similar to locating a patch pipette used in electrophysiological experiments, the shadow of the needle can be used to estimate its location (See **Supplementary Video 1**).

18 Mark the ROI using the microdissection needle. It is not required to completely cut out the tissue, but the cuts should be deep enough so that they are visible under a binocular microscope (See **Supplementary Video 1**).Note that the dissection needle tip isn’t used to cut tissue; Rather the flexibility of the needle allows it to bend into the tissue, pushing into it and hence cutting it. By adjusting the length and thickness of the needle as well as the angle at which the needle approaches the slice, the cut length can be controlled (**Figure 2D** and **E**).

19 Gently remove the slice anchor and transfer the slice to a dissection mat placed under the dissection (binocular) microscope. Ensure that the slice does not fold onto itself and that the microincisions are facing upwards. Remove excess ACSF (**Figure 3**). Optionally, rinse the slice with PBS in order to wash away contaminants and remove the excess PBS.

**Figure 3.**
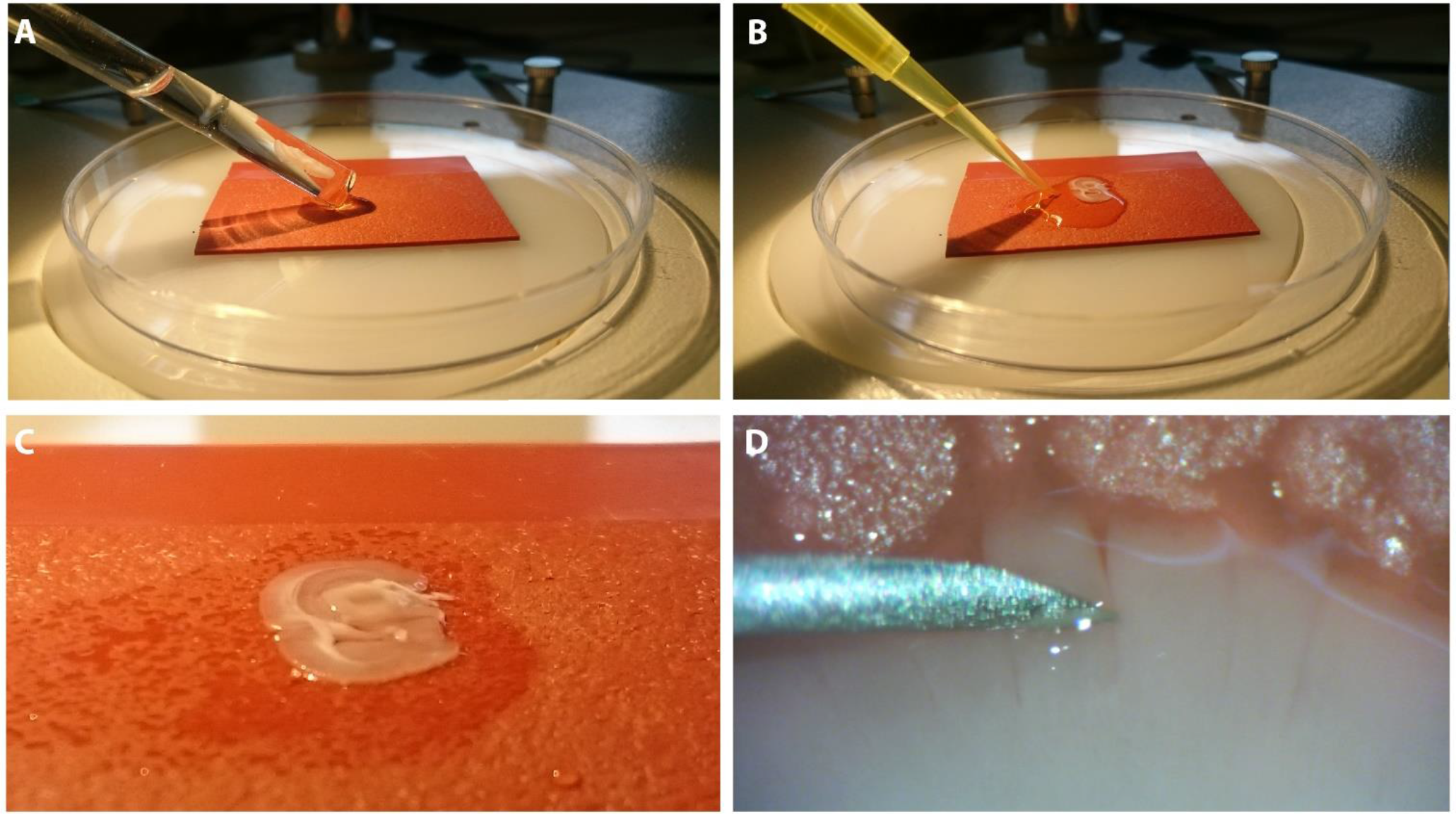
Tissue collection following microdissection. **(A)** Microdissected acute brain slice is transferred from the holding chamber to a flexible dissection mat; care is taken to maintain the same orientation so that previously applied cuts are facing upward to facilitate identification. **(B)** Excess ACSF is removed by pipetting, taking care to reduce mechanical interaction with the slice. Optionally the slice can be rinsed with PBS at this stage. **(C)** Slice, after removing excess medium and optional washing, is now ready for tissue collection. **(D)** A sharp, sterile and clean 32G insulin needle is used for final tissue dissection and collection. Cortical L4 displays increased opaqueness in comparison to L2/3; Combined with its known cortical depth, the two layers can be separated. An additional horizontal microincision can be used if needed. Tissues are collected in Eppendorf tubes and immediately snap-frozen, preserving RNA and protein integrity.

***CRITICAL*** Removing ACSF/PBS will ensure that the slice does not move during tissue dissection and collection. It also helps with cutting out the appropriate tissue as the light reflection from the liquid surface is eliminated.

#### TROUBLESHOOTING

20 Quickly cut out the ROI using an insulin needle (32G). After cutting out the tissue, transfer it to a sterile (RNAse-free) Eppendorf tube using the needle.

***CRITICAL*** Because most of the ACSF is removed, the tissue is very vulnerable at this stage. Additionally, the tissue has been damaged during microdissection, activating cellular response pathways that could alter the tissue’s molecular make-up. Therefore this step should be completed as quickly as possible.

21 Close the tube and immediately snap-freeze it. When completely frozen, the tube can be transferred to dry ice to free up space in the nitrogen container.

23 Store tissue samples at −80 degrees Celsius. Until further use.

***PAUSE POINT*** Samples can be stored indefinitely at −80 degrees Celsius

### RNA isolation

***TIMING*** Approximately 1 - 1.5 hours depending on the number of samples and the isolation method

#### TROUBLESHOOTING

Although tissues obtained using the described technique can be potentially used for any -omics profiling, transcriptomic and proteomic screening are the most commonly used approaches. In the case of the latter, the isolated sample can be thawed and lysed for mass spectroscopic analysis without further tissue manipulation; see Kole et al 2017b for a detailed liquid chromatography mass spectroscopy analysis of samples acquired as described herein.

RNA-sequencing requires RNA isolation. Here we exemplify the RNA isolation pipeline using cortical columns and layers isolated from the rodent somatosensory cortex (see Kole et al 2017a). From each animal, tissues from four slices were obtained. We pooled tissue samples from the same cortical layer and column (e.g. from 4 slices we obtained L2/3 of the C cortical column, and these were pooled at the RNA isolation step. See **Figure 2B** and **C**).

24 Take the first sample to be pooled from the dry ice, and immediately add lysis reagent (e.g. QIAzol). Pipette tissue up and down quickly while taking care not to create bubbles. In our hands, complete lysis of small tissue samples (~300 - 400 μm^3^) is achieved within 15 seconds; larger tissues may have to be crushed before adding lysis reagent.

*CAUTION* some lysis reagents, such as Trizol or QIAzol, are harmful. Take the necessary precautions (work under a fume hood and dispose waste in accordance to local/national/international regulations).

***CRITICAL*** complete and rapid lysis of tissue is critical to obtain high quality RNA

25 Take the second sample to be pooled from the dry ice. Immediately transfer the entire volume from the first tube to the second and repeat the previous step. Once all tissues of a single experimental condition (e.g. all L2/3 samples from column C) have been pooled, place tube on ice and continue with the next tissues (e.g. all L4 samples from column C).

26 When all tissues have been pooled and lysed, continue with the protocol following the manufacturer's instructions. If the protocol includes an on-column DNAse step, it is recommended to perform it; Otherwise perform DNAse treatment after elution and purify the sample.

27 Typically, the final step entails RNA elution from a spin column. In our experience RNA yield can be enhanced by 1) heating the RNAse-free water to 70 degrees Celsius before adding it to the spin column, 2) incubating the water on the spin column for ~1 minute and 3) reusing the eluate, loading it on the same spin column a second time. After elution and in downstream applications, keep RNA samples on ice at all times.

29 Determine RNA quantity using Qubit (<40 ng/μl expected concentration) or Nanodrop (>40 ng/μl expected concentration). Confirm RNA integrity using BioAnalzyer or TapeStation using the kits appropriate for the determined RNA concentration; see *Materials - Equipment* section (**Figure A** and **B**).

30 Store RNA samples at -80 degrees Celsius.

***PAUSE POINT*** Samples can be stored indefinitely at −80 degrees Celsius

#### TROUBLESHOOTING

| Step | Problem | Possible reasons | Solution |
| --- | --- | --- | --- |
| 19 - 20 | Incisions not visible after microdissection | Incisions are not deep enough | Prepare more rigid microdissection needles. Pull shorter pipettes or cut the needle further from the tip (while ensuring that the tip is still thin enough). Alternatively, use a higher magnification of the binocular microscope. |
|  |  | Tissue clings together | Add a small amount of ACSF and use the insulin needle to gently pull at the tissue at the sides of the ROI, this might reveal the cuts. Avoid contact with the tissue of interest. |
|  |  | Slice is the wrong way up | Add ACSF and carefully turn the slice over. Cuts made using microdissection needles are best identified under the binocular microscope if the slice is oriented in the same way as when the microdissection needle was used. |
| 29 | Low RNA or protein integrity | RNA and protein degradation due to poor slice viability | Ensure media are prepared fresh, continuously carbogenated and kept at the indicated temperatures. Work quickly but carefully during slice preparation to avoid tissue damage. Use higher magnification (40x) to assess number of dead or dying cells. Use alternative slice thicknesses according to the animal's age. |
|  |  | RNA and protein degradation due to intermittent thawing of tissue | Before further processing, keep isolated tissues on dry ice, at -80 degrees Celsius or in liquid nitrogen at all times. |
|  |  | RNA degradation due to RNase contamination | During tissue dissection and especially RNA isolation, change gloves regularly. Clean gloves and equipment using RNaseZap. Work quickly during tissue dissection and RNA isolation. |

## Anticipated results

Electrophysiological viability of the slices can be studied, for example, using standard current-clamp and voltage-clamp whole cell-recording protocols to visualize the spiking dynamics and voltage-dependent conductances in the neurons of interest (**Figure 4**). We have performed 450+ such electrophysiological experiments (Lantyer et al (in submission)) that showed that the slice preparation protocol described herein allows neuronal survival for >5h after the Step 14 of the protocol (see **Figure 1**), although accelerated neuron death is observed during the last two hours. Because tissue dissection takes <5 minutes to complete after Step 14 of the protocol, the slices remain viable for the entire sample isolation protocol.

**Figure 4.**
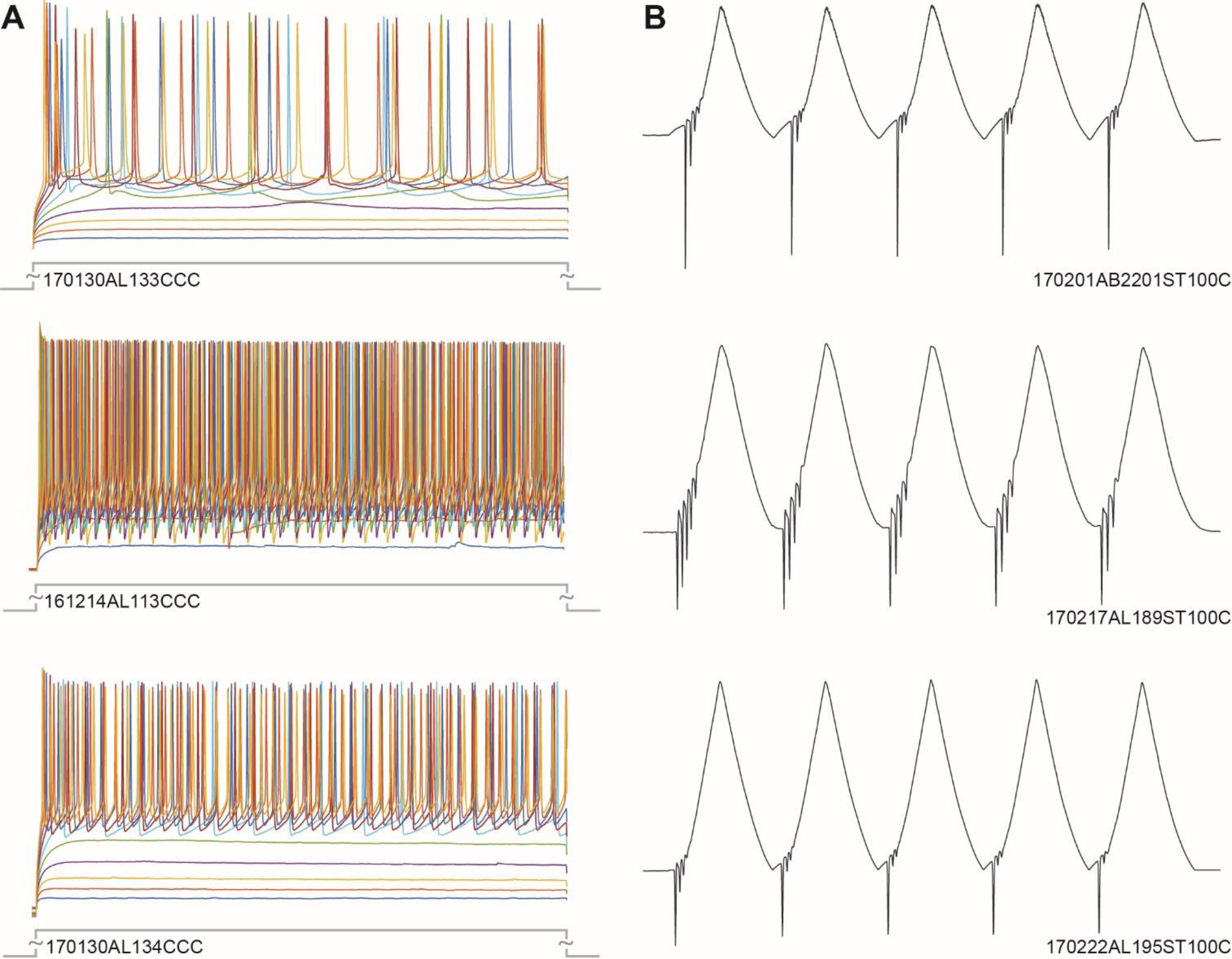
Electrophysiological viability of the neurons. Representative examples of neurons recorded from the supragranular laminae (L2/3) in the barrel cortex after slice preparation as described above. Neurons were recorded 30 min-3h after Step 14 of the protocol in Figure 1. **(A)** Spiking response to step-and-hold somatic current injection. Colors denote varying, cumulative, levels of current amplitude (20 pA/step). **(B)** Voltage-dependent conductances using a saw-tooth voltage clamp protocol (at 5 Hz, 200 ms interpeak interval; peak: +80 mV). The numbers in figurines denote experiment IDs. Raw data from these experiments and 450+ others collected under similar conditions are publically available (see Angelica da Silva Lantyer, Ate Bijlsma, Niccolo Calcini, Fleur Zeldenrust, Wim J. J. Scheenen, TC (in submission)).

For transcriptomic analysis RNA integrity is a key determinant of the outcome of downstream analyses, and should therefore be quantified before sequencing. Well-established methods for quantitative analysis of RNA integrity include the use of Agilent BioAnalyzer and TapeStation, which assess the presence of ribosomal RNA bands, the ratio of their intensities and the presence of smearing (which is an indicator of RNA degradation). In our experience, RNA isolated using the method described herein has a RIN value of >7 (**Figure 5A**) which exceeds the requirements for high-quality RNA-sequencing (see Kole et al, 2017a).

**Figure 5.**
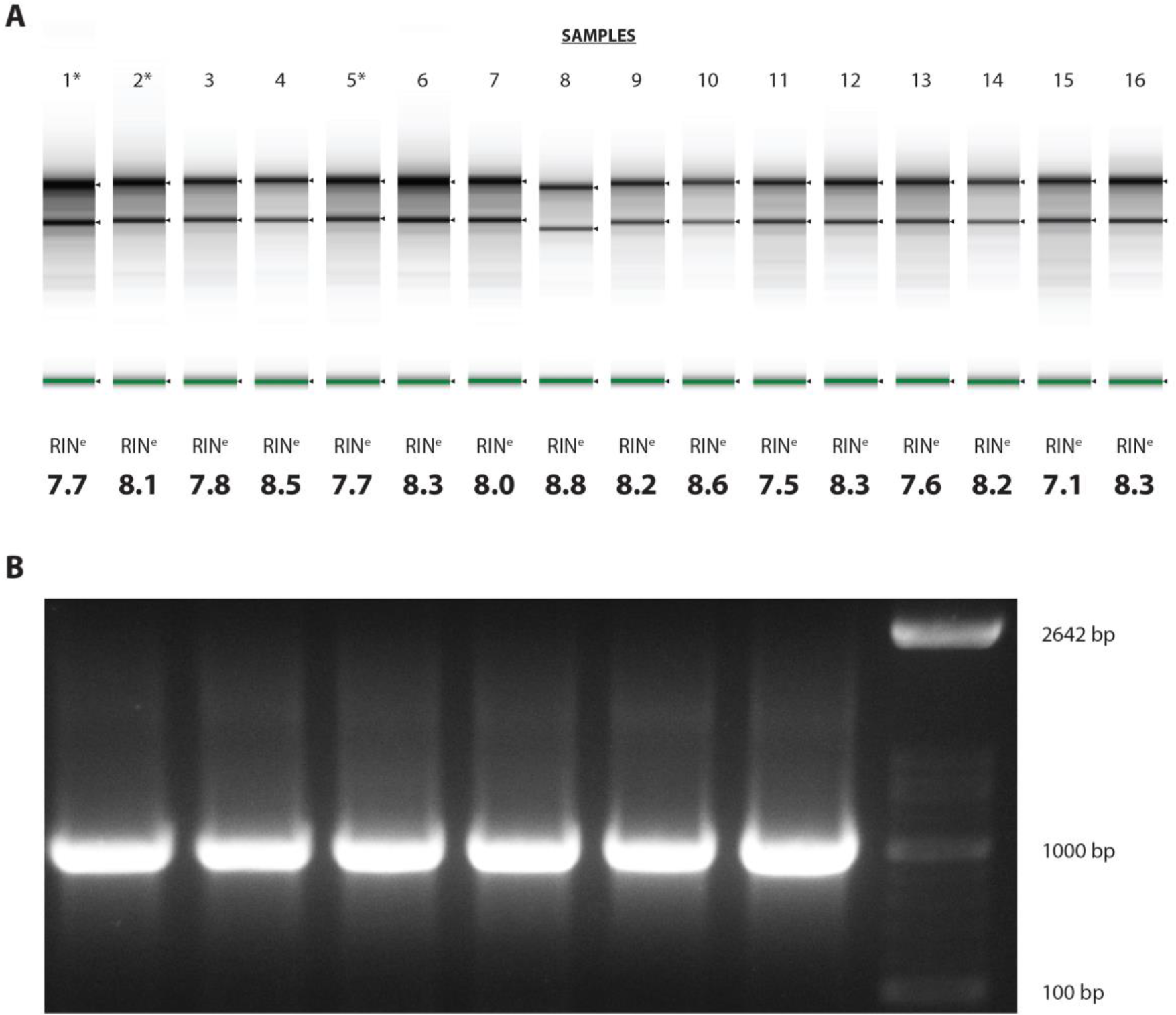
Quality control of total RNA samples. **(A)** RNA integrity as determined by Agilent Tapestation. RNA Integrity Number (RIN) or RIN equivalent (RIN^e^) values of 7 or above are commonly required for successful deep sequencing. Minute samples obtained using the method described readily meet this standard. Asterisks denote samples that are also shown in **B**. **(B)** PCR products using low input RNA samples. Obtained RNA should be free of contaminants in order to be used for cDNA synthesis, the first step in downstream PCR amplification as well as RNA sequencing.

Depending on the expected RNA yield, a Nanodrop or Qubit can be used for total RNA quantification. In our experience, Nanodrop readings become less reliable if concentrations are below ~40 ng/μl. Therefore, if expected concentrations are lower, Qubit (or a similarly sensitive method) might be the preferred choice. Nanodrop is also used to assess the purity of the sample, but also these readings become increasingly less reliable as RNA quantity is reduced. An alternative way to confirm that samples are of sufficient purity is to use it for cDNA synthesis (which is also the first step in RNA-sequencing) and subsequent PCR amplification. If a fairly large amplicon (500 - 1000 bp) can be produced, this indicates that the RNA is suitable for enzymatic reactions (**Figure 5B**).

Many brain regions can be molecularly identified through the expression specific genes ^36^. In the neocortex, for example, laminae differ from one another in their RNA expression profile ^50^ (**Figure 6A**, **Figure 7**). Populations of cell classes also differ between cortical layers: whereas vasoactive intestinal peptide (VIP), vesicular glutamate transporter 1 (VGlut1), calbindin and calretinin-expressing cells are more abundant in L2/3 ^51–53^, parvalbumin, GAD67, NeuN-expressing neurons and oligodendrocytes (distinct in their expression of myelin oligodendrocyte glycoprotein, MOG, and myelin binding protein, MBP) are found at higher densities in L4 ^4,51,54^. The abundance of proteins specific to these cells can be used to confirm laminar specificity of used tissue samples both for transcriptomic and proteomic mapping (**Figure 6B**). This principle can also be reversed: brain regions neighbouring the relevant region but not of interest to the study at hand might be recognized by molecular markers. The (relative) absence of these could help to confirm sufficient precision of tissue extraction.

**Figure 6.**
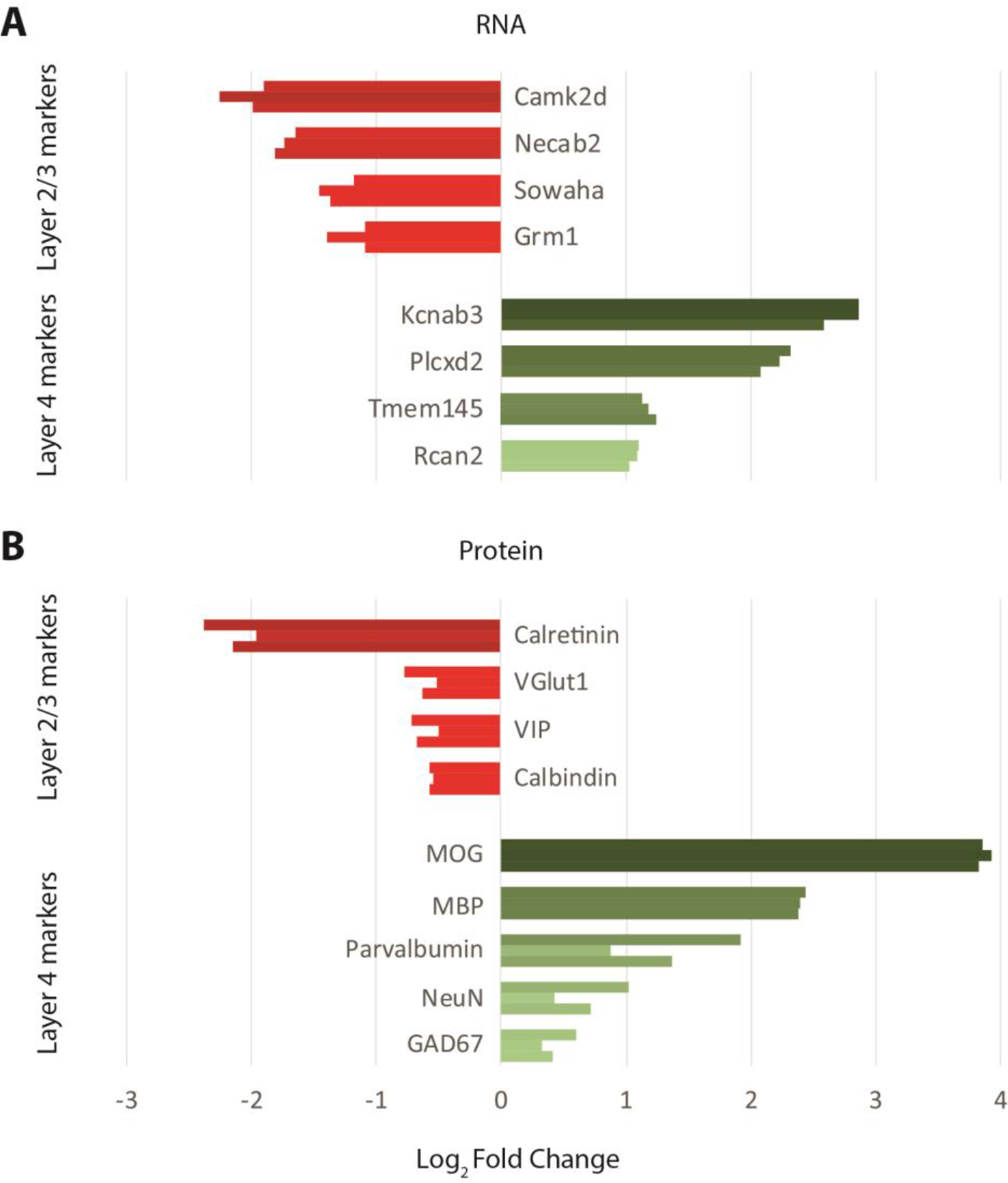
Differential gene transcription and translation between cortical laminae. **(A)** Transcripts (also see **Supplemental Figure 1**) and **(B)** proteins that are preferentially expressed in either supragranular or granular layers (i.e. L2/3 or L4, respectively) follow similar expression patterns in RNA-seq and LC-MS data, confirming laminar specificity of input samples.

**Figure 7.**
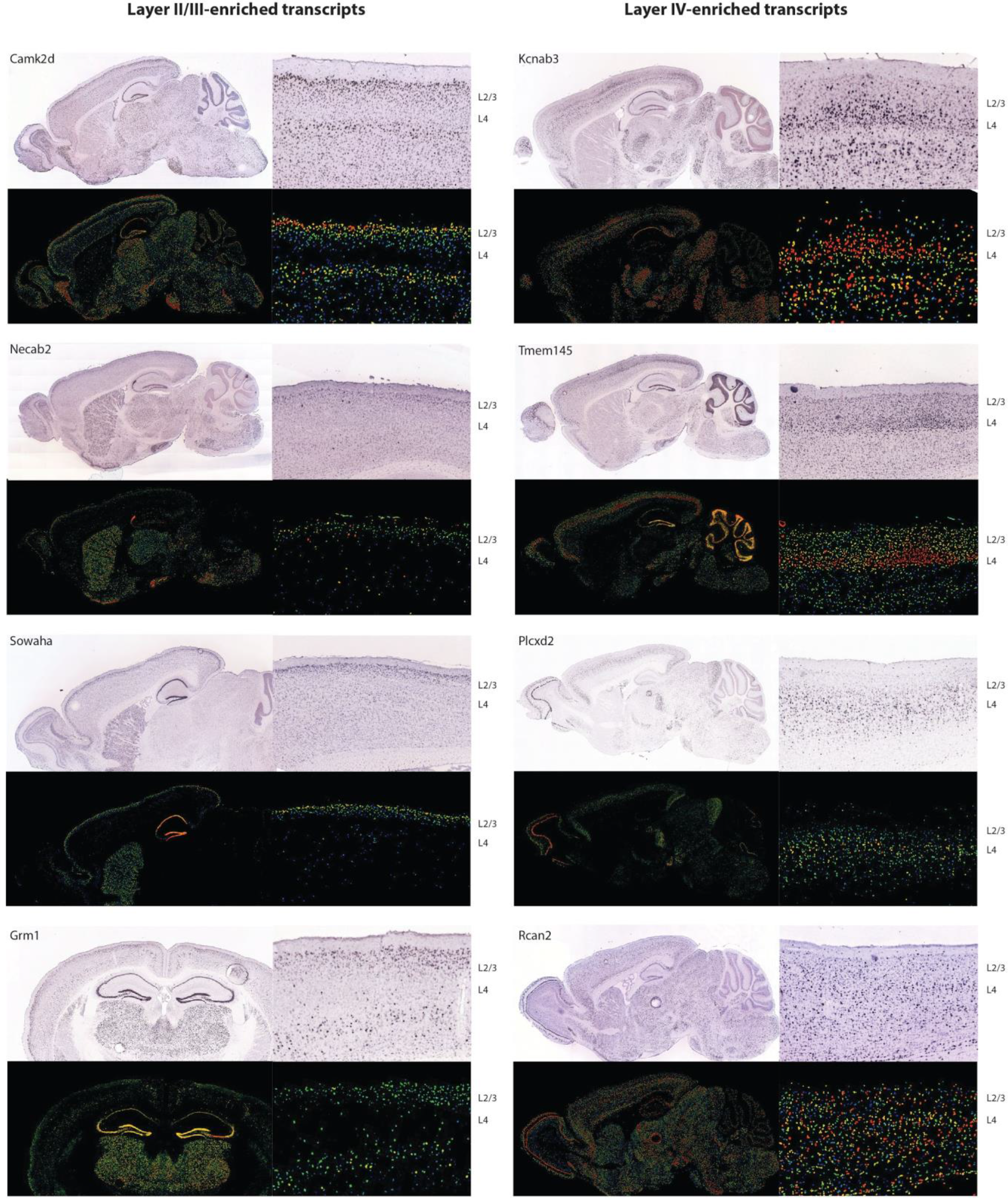
Laminar-specific expression of selected transcripts. Transcripts listed in **Figure 5A** display laminar preference in their expression, as shown by publically available in situ hybridization database (http://mouse.brain-map.org/) from the Allen Brain Institute. For each transcript, top figurines show *in situ* hybridization data; bottom figurines show background subtracted, normalized density of gene transcription ^50^.

---

**Supplemental Video 1. Microdissection of cortical columns and layers of the mouse barrel cortex.** A microdissection needle is used to create intercolumnar incisions to allow identification of individual cortical columns under a dissection microscope (see Supplemental Video 2). The bottom of the incisions indicate the end of L4, allowing laminar separation during tissue collection. Note that the fourth incision required additional depth to ensure ease of visualization, which could be applied readily and with a high degree of precision.

**Supplemental Video 2. Tissue collection of cortical tissues after microdissection.** After microdissection under an upright microscope, slices are transferred to a (binocular) dissection microscope that allows easy access to the tissue. From each column, L2/3 and L4 are separately collected and immediately snap-frozen.

